# The antimicrobial peptides pipeline: a bacteria-centric AMP predictor

**DOI:** 10.1101/2024.05.26.595993

**Authors:** Werner Pieter Veldsman, Qi Zhang, Qian Zhao, Eric Lu Zhang

**Affiliations:** Department of Computer Science, Hong Kong Baptist University, Kowloon, Hong Kong SAR, China; State Key Laboratory of Chemical Biology and Drug Discovery, Department of Applied Biology and Chemical Technology, The Hong Kong Polytechnic University, Hong Kong SAR, China; Centre for Eye and Vision Research, Hong Kong SAR, China; Institute for Research and Continuing Education, Hong Kong Baptist University, Shenzhen, China

## Abstract

Antimicrobial peptides (AMPs), unlike antibiotics, are encoded in genomes. AMPs are exported from the cell after expression and translation. In the case of bacteria, the exported peptides target other microbes to give the producing bacterium a competitive edge. While AMPs are sought after for their similar antimicrobial activity to traditional antibiotics, it is difficult to predict which combinations of amino acids will confer antimicrobial activity. Many computer algorithms have been designed to predict whether a sequence of amino acids will exhibit antimicrobial activity, but the vast majority of validated AMPs in databases are still of eukaryotic origin. This defies common sense since the vast majority of life on earth is prokaryotic. The antimicrobial peptides pipeline, presented here, is a bacteria-centric AMP predictor that predicts AMPs by taking design inspiration from the sequence properties of bacterial genomes with the intention to improve detection of naturally occurring bacterial AMPs. The pipeline integrates multiple concepts of comparative biology to search for candidate AMPs at the primary, secondary and tertiary peptide structure level. Results showed that the antimicrobial peptides pipeline identifies known AMPs that are missed by state-of-the-art AMP predictors, and that the pipeline yields more AMP candidates from real bacterial genomes than from fake genomes, with the rate of AMP detection being significantly higher in the genomes of seven nosocomial pathogens than in the fake genomes.

## Background

Antimicrobial peptides (AMPs) serve a similar purpose to antibiotics, but their chemical structure and *in situ* production differ. Antibiotics are typically monomer products of enzymatic metabolism, while AMPs are polymer products of ribosomal translation. The mechanism of action of AMPs is not limited to cell wall disruption, but extends to intracellular targeting and the elicitation of host immune responses. This versatile mechanism of action confers broad-spectrum antimicrobial activity, making AMPs promising alternatives to traditional antibiotics.

Approximately 11% of the validated AMPs in the Antimicrobial Peptide Database (APD) [1] are of bacterial origin (calculated using values from [2]), with validated AMPs of both plants and animals exceeding that of bacteria. Since prokaryotes are taxonomically more diverse than eukaryotes, the disproportionately low number of validated bacterial AMPs in the APD points to a trove of undiscovered bacterial AMPs. The reason for the underrepresentation of bacterial AMPs in the APD may relate to AMP prediction software often focusing on single machine learning algorithms [3] that exclusively rely on training data from established eukaryotic-biased AMP databases. Moreover, ambiguous biological phenomena such as sequence complexity [4], amino acid periodicity [5,6], and codon usage [7], lead to differences in analysis results between taxonomic kingdoms and peptide structural dimensions (e.g. primary vs secondary vs tertiary peptide structure). These differences can easily be neglected when using datasets from databases that cater for a variety of taxonomic and structural information.

On the other hand, it is essential that features selected for training an AMP predictor are as diverse as possible due to the high heterogeneity of AMPs. Indeed, a tendency in AMP prediction studies to use similar negative data sampling techniques has been shown to significantly bias AMP prediction models to the extent that it affects the validity of reported benchmarking results [8]. The heterogeneity of AMPs demands prediction approaches beyond the standard curriculum. A noteworthy study did bypass reliance on comparative biology by drawing an analogy between a game of cards and AMP prediction [9], however, a bacteria-centric AMP predictor that takes advantage of recent discoveries in protein folding prediction [10] and homology determination [11], is still lacking.

The purpose of the antimicrobial peptides pipeline is to generalize the AMP prediction approach rather than to generalize a ML model. To this end, the pipeline employs numerous methods that each have their own merit grounded in molecular biology principals rather than statistical theory. Assessment results using established AMP predictors show that the antimicrobial peptides pipeline is able to detect known AMPs that fall outside of a 91.6% detection rate of an AMP predictor rated best in a comprehensive assessment of ML based AMP prediction software [3]. The software description that follows describes the antimicrobial peptides pipeline in chronological order as it is executed from start to finish.

## Software description

The antimicrobial peptides pipeline integrates scripts written in R v4.2.0 [12], python v3.12.3 and bash v5.2.21. Execution takes place in a conda environment with bash being the controlling interpreter. The pipeline produces ten batches of AMP candidates using ten distinct methods (Figure 1).

**Figure 1:**
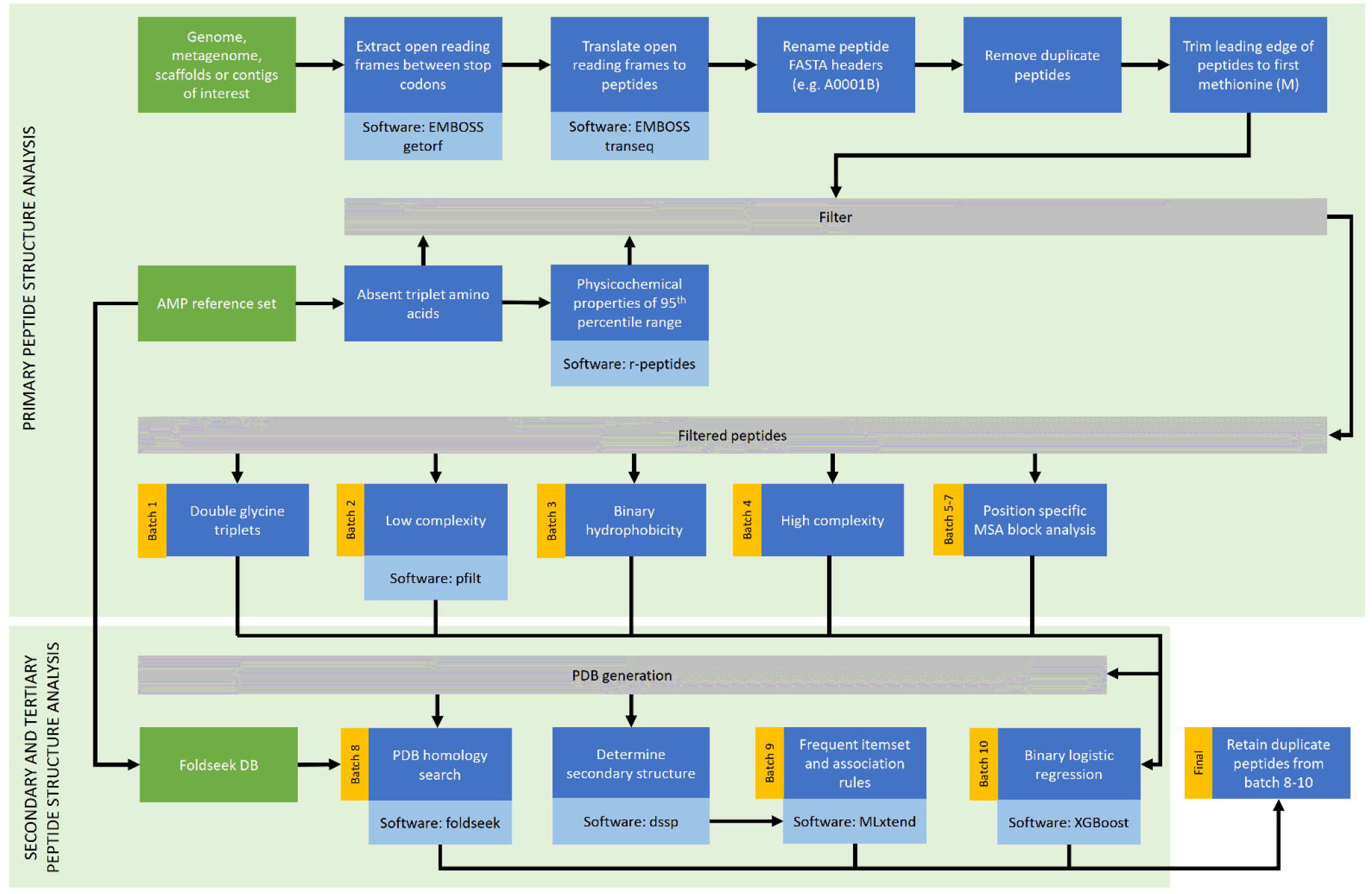
Pipeline flow diagram. The pipeline starts with nucleotide sequences as input (top left) and ends with the creation of final set of AMP candidates (bottom right). Steps that produce AMP candidate batches during execution of the pipeline are shown with yellow sidebars.

The pipeline starts with a single end-to-end command that takes as its only argument a FASTA file with nucleotide sequences. Pipeline parameters can however be tweaked by editing the parameters file located within the project root directory. The first steps in the pipeline invoke prechecks for the existence of software dependencies, the existence of previous results that may be overwritten, and a basic validation of the input file’s FASTA format. After prechecks are completed, the getorf library of EMBOSS v6.6.0.0 [13] is used to extract stop-codon to stop-codon small open reading frames (ORFs) of up to 150 bp in length from the user provided nucleotide sequences. The transeq library of EMBOSS is used to translate the ORFs to peptides using a bacterial codon table. The peptides are then renamed using an alpha-numeric naming scheme for convenient handling of downstream operations, and deduplicated using seqkit v2.8.0 [14]. To obtain AMP candidates that would potentially represent translated peptides in bacterial vectors, peptides are filtered for those that contain at least one methionine residue. The peptides are then trimmed up to the first methionine residue and further filtered to retain only peptides with a length of at least ten residues.

Despite AMP databases being biased in favor of eukaryotes, a biological reference remains useful to constrain AMP candidates using the sequence and physicochemical properties of validated AMPs. The Antimicrobial Peptide Database (APD) is a well-known database of validated AMPs [1]. Tripeptide analysis of the APD revealed 553 combinations of the 20 standard amino acids that are absent in validated AMPs. For this reason, all peptide candidates of the pipeline are filtered to retain only those that do not contain any of the 553 tripeptides. Amino acids in a peptide can be assigned to nine physicochemical categories: tiny, small, aliphatic, aromatic, non-polar, polar, charged, basic, and acidic (as discussed and with software provided in [15]). When each amino acid in a given peptide is classified using these categories, count values arise. An analysis of the count values for the APD reference set was used to determine inter-percentile value ranges (2.5^th^ to 97.5^th^) for each of the nine categories, which are used in the pipeline to further filter AMP candidates. Once inter-percentile range filtering completes, the pipeline is ready to produce the first seven batches of AMP candidates.

The first batch of AMPs is produced by retaining those peptides that contain the double glycine triplet motif GGG[^G]{1,}GGG. In the APD reference set, this doublet glycine triplet motif appears more frequently than all other double triplet motifs of the 19 remaining standard amino acids put together. The second batch is produced by using pfilt v1.5 [16] with default settings to detect low complexity peptides including peptides with coiled-coil, transmembrane and WD repeat signatures. Binary patterns sometimes play a more important role than actual amino acid composition in determining secondary structure, especially when determining the periodicity of peptide features [17]. The third batch therefore consists of peptides containing the HXXHHXXHHX motif after binary hydrophobicity conversion, where the amino acid residues A, I, L, M, F, W & V are represented by H, the amino acid residues S, T, C, N, Q & Y are represented by P, and all others by X. The HXXHHXXHHX motif was selected since it is the most frequent 10-mer at the k-peptide frequency peak of the APD reference set. The fourth batch consists of high diversity peptides that contain at least one of each of the twenty standard amino acid residues. The second and fourth batches are thus complementary in that they represent peptide sequences that are respectively highly conserved [4] and less likely to be conserved. Batches five, six and seven all produce candidate AMPs using motifs that were determined using MAFFT v7.505 [18] and TrimAl v1.4 [19] for multiple sequence alignment followed by compositional data analysis (CODA) for motif discovery (Table 1). Motifs were arbitrarily selected with the aim to assign motif space where a relatively small number of amino acids represented a relatively large proportion of amino acids in the alignment while simultaneously avoiding assignment of motif space when there were no representative amino acids for the majority of the alignment. Two sets were analyzed during motif discovery: (i) validated APD AMPs, and (ii) a set of identical protein group prokaryotic RefSeq sequences downloaded from the NCBI on 13 March 2024 with the search query “antimicrobial peptide”.

**Table 1:**
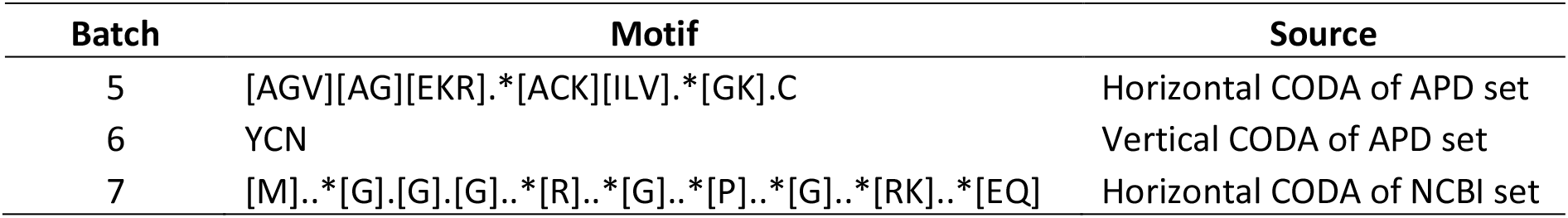
Motifs used for AMP discovery.

Peptides picked up by the first seven methods are collectively referred to as the primary candidate set. The primary candidate set is next used to obtain tertiary structure predictions (in PDB file format) from the evolutionary scale modeling (ESM) Metagenomic Structure Atlas [20] for each peptide sequence in the primary candidate set. The PDB files are used in the production of the eighth and ninth batch. Batch 8 candidates are those whose PDBs pass a homology threshold (length=10; identity=0.5) when queried against a built-in Foldseek [11] PDB database of the APD reference set. Secondary structures such as alpha helices (3-10 and pi), beta sheets (bridged and extended), hydrogen bond turns and loops are determined using DSSP [6,21] before production of the ninth batch. Once secondary structures have been determined, their frequent item set patterns are determined (support=0.1; confidence=0.8) using MLxtend [22], and then compliance filtered using a frequent item set template derived from the APD reference set. The tenth batch is produced using a binary logistic regression model trained by XGboost [23] on eight Moreau-Broto peptide autocorrelation descriptors [24] to predict AMPs and non-AMPs (threshold=0.05). The candidates called by the eighth to tenth batch are collectively referred to as the secondary candidate set. Candidates that appear at least twice in the secondary candidate set are guaranteed to have been called by at least three of the ten methods of the pipeline. These duplicate candidates are referred to as the tertiary candidate set or final candidates. The pipeline ends with a check for the presence of all intermediary files, and the generation of CSV and HTML reports.

## Pipeline assessment

The performance of the pipeline was assessed using AmpGram [25] and amPEPpy [26]. AmpGram outputs a score indicating the probability of a peptide being an AMP, while amPEPpy outputs a binary AMP/non-AMP classification. Both these tools were included in a recent comprehensive assessment of AMP predictors [3] with amPEPpy noted by the authors for having the best predictive performance on an independent data set. Ampgram was used to assign probability scores for (i) the APD set, (ii) AMPs called by the pipeline on the APD set, (iii) AMPs called by the pipeline on a concatenation of 27 real bacterial genomes that are known to produce AMPs [27], (iv) AMPs called by the pipeline on a concatenation of 27 fake genomes with randomized nucleotides, (v) ORFs extracted by the pipeline from the same 27 real genomes, and (vi) ORFs extracted by the pipeline from the same 27 fake genomes. The first two probability result sets are shown on Figure 2 as positive controls with their expectedly higher areas under the curve (AUCs), while the remaining four assessments are tests of the pipeline on the complete genomes of known AMP producers that were downloaded from the NCBI on 22 April 2024 (Table S1).

**Figure 2:**
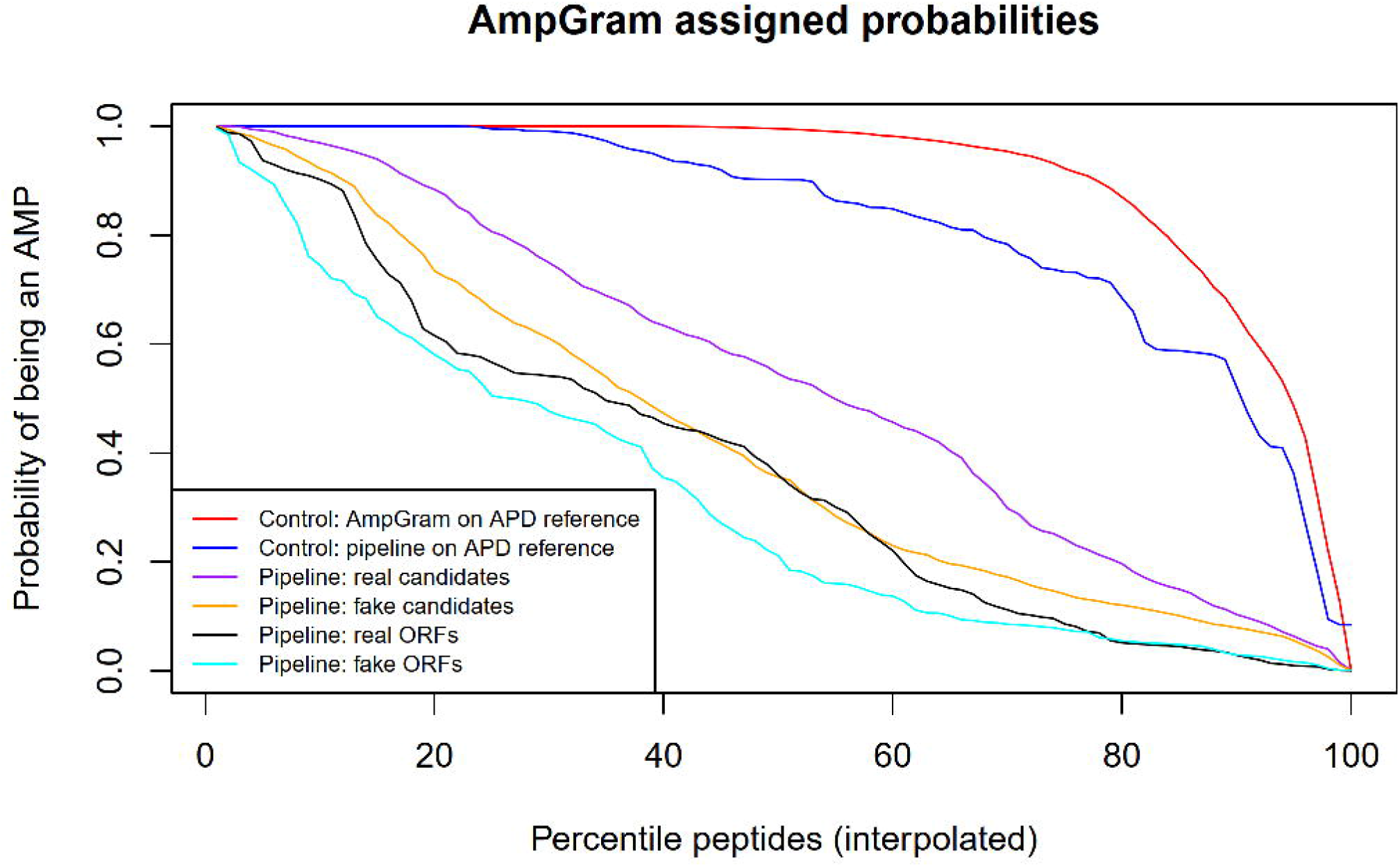
AMP probability scoring. The pipeline was assessed by AmpGram [25] probability scoring of its output. The pipeline has a different prediction profile than AmpGram as shown by the controls, while AMP candidates and extracted ORFs from real genomes are more likely to have antimicrobial activity than AMP candidates and extracted ORFs from fake genomes.

The two controls, where ideally AUC=1, show that the pipeline has a different AMP candidate selection strategy than AmpGram (AUC=0.906 vs AUC=0.819). Evaluation of the pipeline on real and fake datasets showed that candidates from real genomes are more likely to be AMPs than candidates from fake genomes that mimic the real genomes in contig and nucleotide numbers (AUC=0.532 vs AUC=0.416). ORFs from real genomes are also more likely to encode AMPs than ORFs from fake genomes (AUC=0.373 vs AUC=0.312).

Assessment of the pipeline using amPEPpy was carried out on (i) the APD set, (ii) primary candidates called by the pipeline on the APD set, (iii) AMPs called by the pipeline on a concatenation of 27 real bacterial genomes that are known to produce AMPs [27], (iv) AMPs called by the pipeline on a concatenation of 27 fake genomes with randomized nucleotides, and (v) AMPs called by the pipeline from the genomes of seven multi-drug resistant nosocomial ESKAPE pathogens (*Enterococcus faecium, Staphylococcus aureus, Klebsiella pneumoniae, Acinetobacter baumannii, Pseudomonas aeruginosa* and *Enterobacter cloacae*) See [28] and [29] for ESKAPE acronym deviations. The assessment of the APD set was carried out by pipeline injection since backtranslation to their nucleotide sequence would introduce nucleotide ambiguity. Results showed that while amPEPpy correctly detects 91.7% of the validated APD AMPs (Table 2), the antimicrobial peptides pipeline tends to select those validated AMPs that are not detected by amPEPpy as indicated by the observed positive prediction rate dropping to 16.3%. The difference in the number of detected AMPs between the 27 real and 27 fake genomes are not significant according to three statistical tests for categorical data (Fisher’s exact test, Chi-squared test, Yates corrected chi-squared test (FCY): all p>0.05). Although the pipeline detected half as much more AMPs in the ESKAPE group than in the 27 real genomes, the difference is also not significant (FCY, all p>0.05). However, the number of detected AMPs in the ESKAPE group is significantly different from the number of detected AMPs in the 27 fake genomes (FCY, all p<0.05).

**Table 2:** AMP predictions called by AmPEPpy.

| Set | AMP |  | Total | Observed positive rate | Expected positive rate | Pipeline derived |
| --- | --- | --- | --- | --- | --- | --- |
|  | Yes | No |  |  |  |  |
| APD AMPs (control 1) | 2900 | 262 | 3162 | 91.7 | 100 | No |
| APD AMPs (control 2) | 17 | 87 | 104 | 16.3 | 100 | Yes |
| 27 real genomes | 52 | 703 | 755 | 6.9 | Unknown | Yes |
| 27 fake genomes | 48 | 859 | 907 | 5.3 | Unknown | Yes |
| ESKAPE genomes | 20 | 169 | 189 | 10.6 | Unknown | Yes |

Given that the two controls relied on exactly the same data, the antimicrobial peptides pipeline is highly complementary to amPEPpy and may support other AMP prediction approaches that are based on a design philosophy that places emphasis on statistical procedure by comparing and benchmarking AMP predictors in terms of which statistical procedure was used (e.g. HMMs, RFs or SVMs) rather than placing emphasis on the biological nature of sequences such as the evolutionary role of sequence complexity, the functional role of sequences that may be masked by residue ambiguity, and the simultaneous consideration of homology at primary, secondary and tertiary levels.

## Summary

The antimicrobial peptides pipeline produces ten batches of AMP candidates with due consideration to biological phenomena such as peptide complexity and residue ambiguity. AMP homology searches are conducted in the primary, secondary and tertiary dimension after physicochemical and k-mer properties of validated AMPs are used to constrain AMP detection. The pipeline also employs motifs discovered by multiple sequence alignment of known AMPs, the frequency of secondary structural features, and a machine learning model trained on biological relevant autocorrelation descriptors. Assessment of the pipeline using state-of-the-art AMP predictors showed that its approach to AMP detection could serve as a complementary tool to AMP predictors based exclusively on machine learning algorithms.

## Data availability

Download and installation instructions: https://github.com/Werner0/antimicrobial_peptides

